# Choice-Related Activity during Visual Slant Discrimination in Macaque CIP but Not V3A

**DOI:** 10.1101/457531

**Authors:** L. Caitlin Elmore, Ari Rosenberg, Gregory C. DeAngelis, Dora E. Angelaki

## Abstract

Creating three-dimensional (3D) representations of the world from two-dimensional retinal images is fundamental to many visual guided behaviors including reaching and grasping. A critical component of this process is determining the 3D orientation of objects. Previous studies have shown that neurons in the caudal intraparietal area (CIP) of the macaque monkey represent 3D planar surface orientation (i.e., slant and tilt). Here we compare the responses of neurons in areas V3A (which is implicated in 3D visual processing and which precedes CIP in the visual hierarchy) and CIP to 3D oriented planar surfaces. We then examine whether activity in these areas correlates with perception during a fine slant discrimination task in which monkeys report if the top of a surface is slanted towards or away from them. Although we find that V3A and CIP neurons show similar sensitivity to planar surface orientation, significant choice-related activity during the slant discrimination task is rare in V3A but prominent in CIP. These results implicate both V3A and CIP in the representation of 3D surface orientation, and suggest a functional dissociation between the areas based on slant-related decision signals.

**Significance Statement:** Surface orientation perception is fundamental to visually guided behaviors such as reaching, grasping, and navigation. Previous studies implicate the caudal intraparietal area (CIP) in the representation of 3D surface orientation. Here we show that responses to 3D oriented planar surfaces are similar in CIP and V3A, which precedes CIP in the cortical hierarchy. However, we also find a qualitative distinction between the two areas: only CIP neurons show robust choice-related activity during a fine visual orientation discrimination task.

## Introduction

Perception of three-dimensional (3D) surface orientation is essential for many visually guided behaviors. Electrophysiological studies have identified 3D orientation selective neurons in multiple brain regions of non-human primates (Murata et al., 2000; Taira et al., 2000; Hinkle and Connor, 2002; Sugihara et al., 2002; Nguyenkim and DeAngelis, 2003; Liu et al., 2004; Durand et al., 2007; Sanada et al., 2012; Alizadeh et al., 2018). In particular, the caudal intraparietal area (CIP) represents all combinations of slant and tilt, two angular variables that specify the 3D orientation of a planar surface Rosenberg et al., 2013). Anatomical as well as functional magnetic resonance imaging data suggest that V3A, which precedes CIP in the visual hierarchy, may also contribute to 3D visual processing (Nakamura et al., 2001; Tsao et al., 2003). V3A neurons have two-dimensional orientation (Zeki, 1978c, b, a) and binocular disparity (Anzai et al., 2011) tuning, but their responses to 3D surface orientation have not been examined. Moreover, few studies have tested for functional correlations between neuronal activity and 3D orientation perception. Previous work indicates that reversible inactivation of CIP results in small but consistent deficits in a 3D curvature detection task (Van Dromme et al., 2016), and may produce a deficit in the ability to perform a delayed match-to-sample task in which planar tilt is coarsely manipulated (Tsutsui et al., 2001).

Here we measured the responses of V3A and CIP neurons to 3D surface orientation, as well as their functional correlations with behavior during a fine slant discrimination task. First, 3D surface orientation tuning was measured during a fixation task. The two areas were found to contain similar proportions of selective neurons, as well as similar degrees of selectivity. Second, neuronal activity was recorded while the monkeys viewed planar surfaces at different slants and reported the slant direction in a two-alternative forced-choice task. Receiver operating characteristic (ROC) analysis was used to quantify neuronal sensitivity and to assess choice-related activity (Celebrini and Newsome, 1994; Britten et al., 1996; Dodd et al., 2001; Nienborg and Cumming, 2006; Gu et al., 2007). In contrast to the similarity of stimulus selectivity in the two areas, significant choice-related activity was rare in V3A but prominent in CIP. To further dissociate the contributions of stimulus and choice to neuronal activity, we performed a partial correlation analysis to assess how much variance in the neuronal activity could be attributed to the stimulus and the choice (Zaidel et al., 2017). This analysis confirmed a similar degree of stimulus-related activity in the two areas, and much stronger choice-related activity in CIP than V3A. These results implicate both V3A and CIP in visual surface orientation processing, and demonstrate that binary decision signals during slant discrimination are carried by the most sensitive CIP (but not V3A) neurons.

## Materials and Methods

### Subjects and surgery

All surgeries and experimental procedures were approved by the Institutional Animal Care and Use Committee, and were in accordance with NIH guidelines. Neuronal recordings were obtained from five hemispheres in three male rhesus monkeys (*Macaca mulatta*), denoted as monkeys N, P, and Z, weighing 4-5 kg at the start of the study. Animals were chronically implanted with a lightweight plastic ring for head restraint, a recording grid, and scleral eye coils for monitoring binocular eye movements (CNC Engineering). After recovery, they were trained using standard operant conditioning procedures to fixate visual targets for fluid reward, and to report the direction of surface slant using eye movement responses. After training, neuronal recordings began. We recorded from CIP in two monkeys (N and P), and from V3A in two monkeys (Z and P). Prior to the study, monkey Z underwent a bilateral labyrinthectomy as part of another project. Results from V3A in monkeys Z and P were compared statistically using Wilcoxon rank-sum tests, and no significant differences were found, indicating that the labyrinthectomy had no detectable effects on the current study. Specifically, there were no significant differences in: median choice probability (monkey Z = 0.50; monkey P = 0.47; *p* = 0.73), neuronal threshold (Z = 31.29; P = 23.73; *p* = 0.60), surface orientation discrimination index (Z = 0.68; P = 0.71; *p* = 0.89), squared choice partial correlation (Z = 0.003; P = 0.01; *p* = 0.20), and squared slant partial correlation (Z = 0.02; P = 0.02; *p* = 0.59). A lack of effects of the labyrinthectomy on visual discrimination is not surprising given that the monkeys were head-fixed during the experiments and that previous studies found that visual heading discrimination performance is largely normal within days following a bilateral labyrinthectomy (Gu et al., 2007).

### Data acquisition

Epoxy-coated tungsten microelectrodes (Frederick Haer Company, diameter 125 μm, impedance 1-5 MΩ at 1kHz) were inserted into the cortex through a transdural guide tube using a hydraulic microdrive to record extracellular action potentials. Neuronal voltage signals were amplified, filtered (1Hz – 10 kHz), and displayed on an oscilloscope to isolate single units using a window discriminator (BAK Electronics). Raw voltage signals were digitized at a rate of 25 kHz using a CED Power 1401, and single units were sorted offline as needed (Spike2; Cambridge Electronic Design). In some experiments, action potentials were displayed and isolated using the SortClient software (Plexon).

The CARET software was used to segment visual areas in magnetic resonance imaging (MRI) scans of monkeys N and P (Lewis and Van Essen, 2000). Recording sites were localized to CIP (which the Lewis and Van Essen Atlas designates as the lateral occipitoparietal zone) using the resulting MRI atlases (Van Essen et al., 2001; Rosenberg et al., 2013). When lowering an electrode dorsal-ventrally, CIP was preceded by either the intraparietal sulcus or by cells with prevalent eye-movement responses, depending on the medial-lateral position of the penetration. Once either the intraparietal sulcus or eye-movement responsive cells were passed, neurons were tested for surface orientation selectivity. Neurons in CIP were further identified as having large receptive fields often extending into the ipsilateral visual hemifield (Taira et al., 2000). Area V3A was targeted using the MRI atlas in monkey P and using stereotaxic coordinates in monkey Z. Area V3A is located ventral-lateral and adjacent to CIP. Lateral to CIP and dorsal to V3A is a large patch of white matter. Thus, both CIP and gray/white matter transitions provided landmarks for targeting V3A. As electrodes were advanced dorsal-ventrally, observed gray/white matter transitions were compared with coronal sections to localize V3A. Receptive field mapping was used to compare the receptive field sizes of V3A neurons to previously published data. Receptive field size increased with eccentricity (*r* = 0.621, *p* = 0.002), and the linear fit y = 0.47x + 1.8 was similar to previous measurements: y = 0.33x + 1.78 (Galletti and Battaglini, 1989) and y = 0.38x + 2.8 (Nakamura and Colby, 2000) obtained using DataThief (Tummers, 2006). We compared response latency between the areas, and found that V3A neurons (median = 56 ms) responded significantly faster than CIP neurons (median = 72 ms), Wilcoxon rank-sum test, *p* = 0.02.

### Behavioral control and stimulus presentation

Behavioral control was carried out with custom Spike2 scripts. The monkeys sat in a primate chair ~ 32.5 cm from a liquid crystal display (LCD) on which stimuli were displayed (System 1: NEC Accusync LCD 93VX; System 2: Dell 1707 FP). An aperture constructed from a black non-reflective material was affixed to the screen such that the monkey could only see stimuli within a 30 cm (System 1) or 18 cm (System 2) diameter circular aperture. The same material extended between the LCD and the monkey, occluding the view of the surrounding room. The OpenGL graphics library was used to program visual stimuli that were generated using an OpenGL accelerator board (Quadro FX 3000G, PNY Technologies). The fixation point (yellow in color) was presented directly in front of the monkey at eye level and screen distance. Fixation was enforced using 2° version and 1° vergence windows. Due to eye coil failures in monkey P, the binocular eye movements of this animal were monitored in all experiments using an infrared optical eye tracker (ISCAN).

### 3D surface orientation tuning

Surface orientation tuning was measured as previously described (Rosenberg et al., 2013; Rosenberg and Angelaki, 2014a, b). Briefly, a planar surface with a checkerboard pattern was used to measure the joint tuning for slant and tilt (Fig. 1A). Stimuli subtended either 50° or 31° of visual angle. Initial recordings with monkey N were conducted in System 1 (used in our previous CIP studies) which allowed us to present 50° stimuli (30 neurons). However, monkey N outgrew the system, which only accommodates relatively small animals. The remaining data for monkey N (14 neurons) and all data from monkeys P and Z were gathered in System 2, for which the largest possible stimulus was 31°. Wilcoxon rank-sum tests revealed no significant differences in the results for monkey N across the two systems, including comparisons of median values of: choice probability (System 1 = 0.57; System 2 = 0.58; *p* = 0.44), neuronal threshold (System 1 = 37.96; System 2 = 31.79; *p* = 0.89), behavioral threshold (System 1 = 3.60; System 2 = 3.74; *p* = 0.27), point of subjective equality (System 1 = 0.16; System 2 = −0.71; *p* = 0.13), squared choice partial correlation (System 1 = 0.02; System 2 = 0.009; *p* = 0.47), and squared slant partial correlation (System 1 = 0.01; System 2 = 0.02; *p* = 0.97).

**Figure 1.**
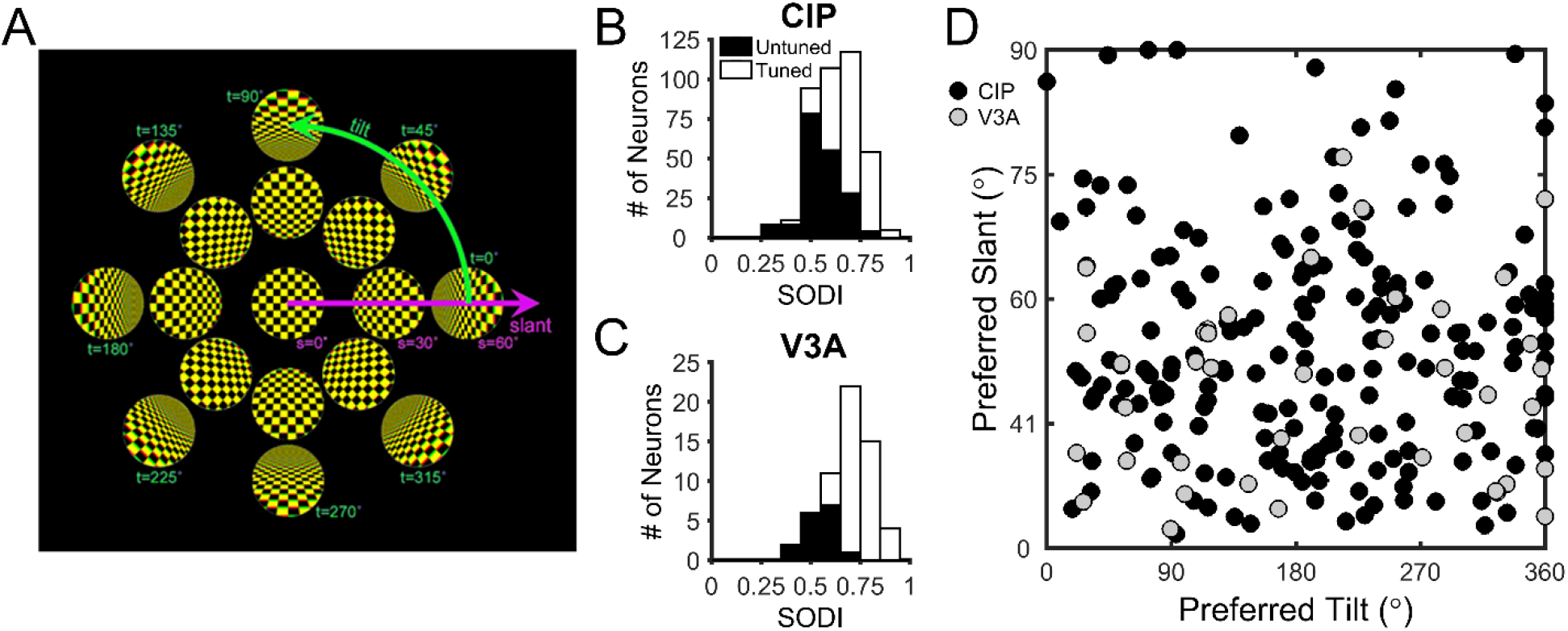
Surface orientation tuning. ***A***, The 3D orientation of a planar surface can be described by two variables, slant and tilt. Tilt specifies the axis within the frontoparallel plane about which the plane is rotated, and slant specifies how much it is rotated. These variables define a polar coordinate system. ***B & C***, Distributions of the surface orientation discrimination index (SODI) for 396 CIP (***B***) and 60 V3A (***C***) neurons. Open bars denote tuned neurons (215 CIP and 44 V3A), and filled bars denote untuned neurons (181 CIP and 16 V3A). ***D***, Equal area projection (Rosenberg et al., 2013) showing joint distribution of preferred slants and tilts for the 215 tuned CIP neurons (black circles) and 44 tuned V3A neurons (gray circles).

Slant was varied between 0° and 60° in 20° steps, and tilt was varied between 0° and 315° in 45° steps. All stimuli were centered on the fixation point and covered the same retinotopic area. Stereoscopic cues were created by rendering the stimuli as red-green anaglyphs. Each trial began with the monkey fixating a point on a blank screen for 300 ms. Fixation was maintained while a checkerboard stimulus was presented for 1,000 ms, followed by 50 ms of fixation with a blank screen. There was a 1,000 ms blank screen inter-trial interval. Stimuli were presented in pseudorandom order. Surface orientation selectivity was assessed for all cells held for at least three repetitions of each stimulus. At most seven repetitions of each stimulus were recorded. For each selective neuron (see Results), a one-way ANOVA was performed to determine if there was significant slant tuning along the 90°/270° tilt axis (see Figs. 1A, 3A,B). Neurons with significant tuning were studied further in the slant discrimination task.

### Slant discrimination task

The slant discrimination task was always performed along the 90°/270° tilt axis. To simplify the description of surface orientation, we do not refer to tilt for the slant discrimination task but instead denote planes with a tilt of 90° (top of the plane closer to the monkey) as having a negative slant, and planes with a tilt of 270° (top of the plane further from the monkey) as having a positive slant (c.f., Rosenberg and Angelaki, 2014b). As illustrated in Fig. 2A, each trial of the slant discrimination task began with the monkey fixating a target on a blank screen for 300 ms after which a random dot stereogram (RDS) depicting a planar surface was presented for 1,000 ms. After presentation of the RDS, the fixation point disappeared, and two choice targets appeared 8.6° above/below the location of the fixation point. The monkey then made an eye movement to one of the choice targets to indicate the perceived slant. Correct responses were defined as a saccade to the upper target when the slant was positive (top-far) or to the lower target when the slant was negative (top-near). Correct responses were rewarded with a drop of water or juice. For planes with slant = 0° (i.e., frontoparallel), responses were rewarded pseudo-randomly 50% of the time. If the monkey broke fixation at any point during the stimulus presentation, the trial was aborted and the data discarded.

**Figure 2.**
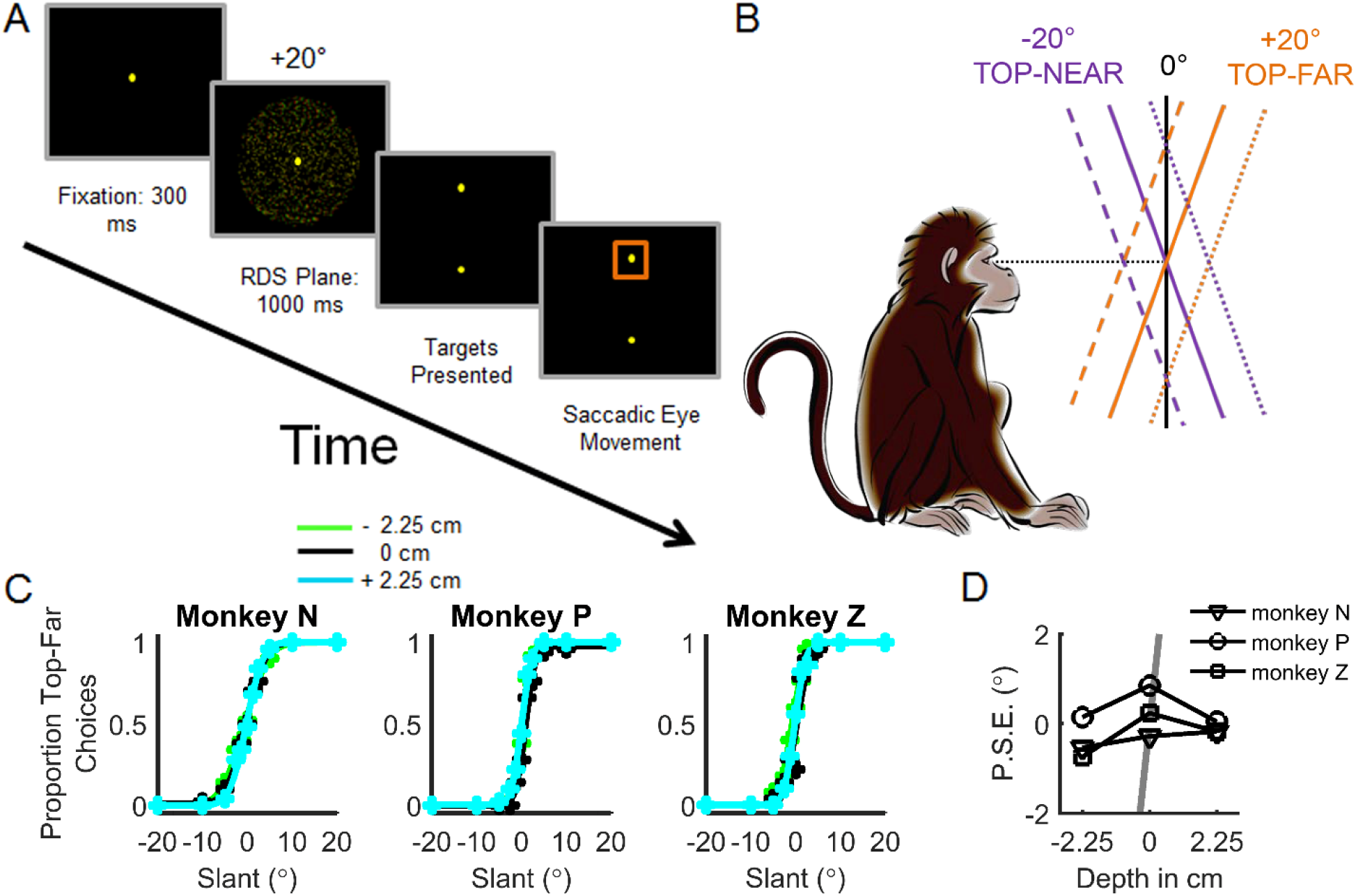
Slant discrimination task and behavioral performance. ***A***, Temporal sequence of events in the slant discrimination task. Each trial began with the presentation of a fixation point at the center of the screen. The monkey fixated this point for 300 ms after which a random dot stereogram (RDS) plane was presented for 1,000 ms while fixation was maintained. The monkey then reported which direction the plane was slanted away from frontoparallel by making a saccade to the upper target if the slant was positive (top-far) or the lower target if the slant was negative (top-near). ***B***, Side view of the task illustrating positive versus negative slants. Solid lines depict planes centered at the fixation depth (screen distance ~ 32.5 cm). Dashed and dotted lines depict planes centered at either near or far depths (2.25 cm in front of or behind the display), respectively. ***C***, Discrimination behavior plotted as the proportion of top-far choices as a function of slant. Data are fit with a cumulative Gaussian for each depth (N = 450 trials/data point). ***D***, The point of subjective equality (P.S.E.) as a function of depth for each monkey. For comparison, the gray line shows the expected dependency of the P.S.E. on stimulus depth if the task was performed based on local absolute disparities rather than slant. The line reaches ±14° at ±2.25 cm but is clipped at ±2° to not obscure the data.

During pilot work, we observed that local orientation cues in checkerboard stimuli could be used to perform the task without having to judge slant. To avoid this potential confound, the discrimination task was performed using RDS planes with uniform dot density on the screen (Sanada et al., 2012). In CIP, slant tuning curves measured with planar surfaces with a checkerboard pattern or a random dot pattern are highly correlated (Rosenberg and Angelaki, 2014b). To discourage the monkeys from using local depth cues to perform the task (Hillis et al., 2004), we varied the mean depth (near = −2.25 cm from the screen, screen distance = 0 cm, far = 2.25 cm from the screen) of the RDS plane from trial to trial (Fig. 2B). This discouraged them from judging whether the upper (lower) half of the stimulus was in front of (behind) the plane of the display. If the animals relied on the absolute disparity of a sub-region of the stimulus to perform the task, large behavioral biases would result at the near/far depths. For the 31° stimulus, biases of at least 14° in magnitude (the slant at which a stimulus would start to cross the screen) would occur in opposite directions for the near and far depths. Behavioral data clearly show this was not the case (Fig. 2C,D), suggesting the animals correctly learned to judge the slant sign. To maintain this behavior during the neural recordings, stimuli were presented at screen distance for 70% of trials, the near depth for 15% of trials, and the far depth for 15%. For the neural recordings, there was sufficient data to reliably analyze the responses measured at screen distance only.

Slant was varied between ±20° with the intermediate slant values tailored to each animal’s performance. For monkeys N and Z, slants of ±20°, 10°, 5°, 2.5°, 1.25°, and 0° were used. For monkey P, slants of ±20°, 9°, 4.05°, 1.83°, 0.82°, and 0° were used. Neurons were recorded while the monkey performed the task for a minimum of 10 repetitions of each stimulus. Sufficient repetitions were recorded for 65 CIP and 23 V3A neurons.

### Data analysis

Analyses were performed in MATLAB (*MathWorks*). The tuning strength of each neuron was evaluated using a surface orientation discrimination index (SODI) following (Prince et al., 2002) and calculated using the full slant–tilt tuning curve. The SODI quantifies the strength of response modulation relative to overall response variability:

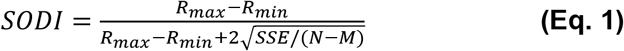

where *R_max_* and *R_min_* are the maximum and minimum responses, respectively. *SSE* denotes the sum squared error around the mean responses, *N* is the total number of trials, and *M* is the number of tested slant–tilt combinations (*M* = 25). Neurons with strong response modulation relative to their variability have SODI values closer to 1, whereas neurons with weak response modulation have SODI values closer to 0.

Behavioral performance in the slant discrimination task was quantified by plotting the proportion of top-far choices as a function of stimulus slant. The resulting psychometric function was fit with a cumulative Gaussian using the Psignifit toolbox (Wichmann and Hill, 2001). The point of subjective equality and behavioral threshold were defined as the mean and standard deviation of the cumulative Gaussian fit, respectively.

Neuronal sensitivity was measured by using ROC analysis to assess the ability of an ideal observer to discriminate between two opposite slants (e.g., −20° from +20°; Fig. 3E,F) based on the firing rate of a recorded neuron and a hypothetical ‘anti-neuron’ with opposite tuning (Britten et al., 1996; Gu et al., 2007). To construct a neurometric function that could be directly compared to the psychometric function (Fig. 4A,B), ROC values were plotted as a function of slant and fit with a cumulative Gaussian using the Psignifit toolbox. Neuronal threshold (an inverse measure of sensitivity) was defined as the standard deviation of the cumulative Gaussian fit. Neuronal and behavioral thresholds were calculated from simultaneously gathered data, allowing for a direct comparison. For this comparison, neuronal thresholds were multiplied by 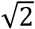 to account for the behavioral task being conducted as a one interval task (Hillis et al., 2004), but the neurometric functions being calculated by comparing two distributions (the neuron and its hypothetical anti-neuron). The time course of neuronal sensitivity was assessed by computing neuronal thresholds in 200 ms time windows shifted every 50 ms over the 1,000 ms stimulus duration.

**Figure 3.**
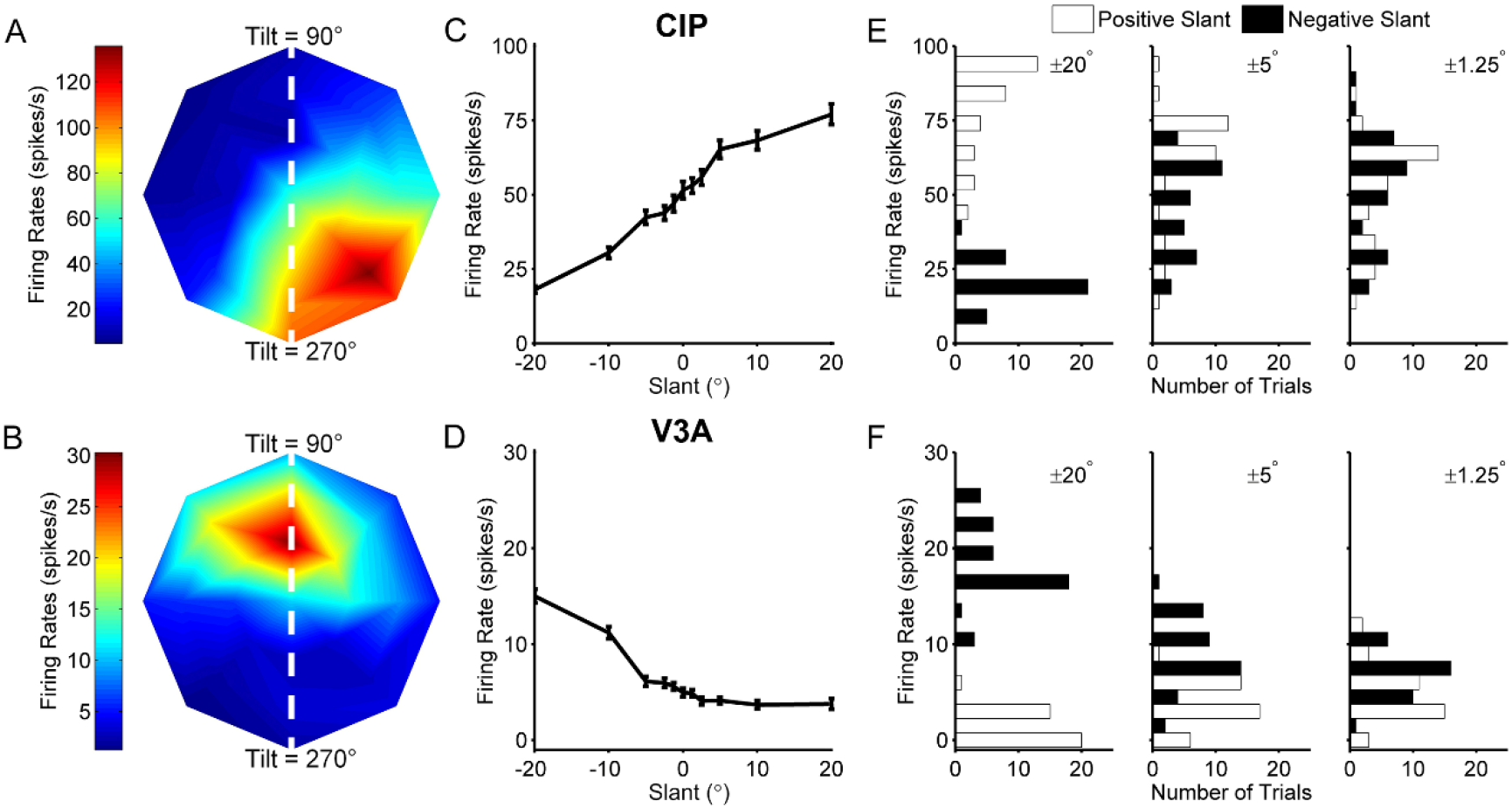
Surface orientation tuning of example CIP and V3A neurons. ***A & B***, Slant–tilt tuning profiles of representative CIP (***A***) and V3A (***B***) neurons. Firing rate is color coded with red hues indicating larger firing rates. The peak of the CIP tuning profile is in the lower right corner, indicating that the cell responded best to a planar surface with the lower right corner closest to the monkey. The peak of the V3A tuning profile is in the upper portion of the plot, indicating that the cell responded best to a planar surface with the top closest to the monkey. White dashed lines correspond to the 90°/270° tilt axis along which the slant discrimination task was performed. *C & D*, Slant tuning curves of the same CIP (***C***) and V3A (***D***) neurons measured during the slant discrimination task. Error bars denote SEM. ***E & F***, Neuronal response distributions for three pairs of slant angles (±20°, ±5°, ±1.25°) for the CIP (***E***) and V3A (***F***) neurons. Negative slants are shown as black bars and positive slants are shown as white bars.

**Figure 4.**
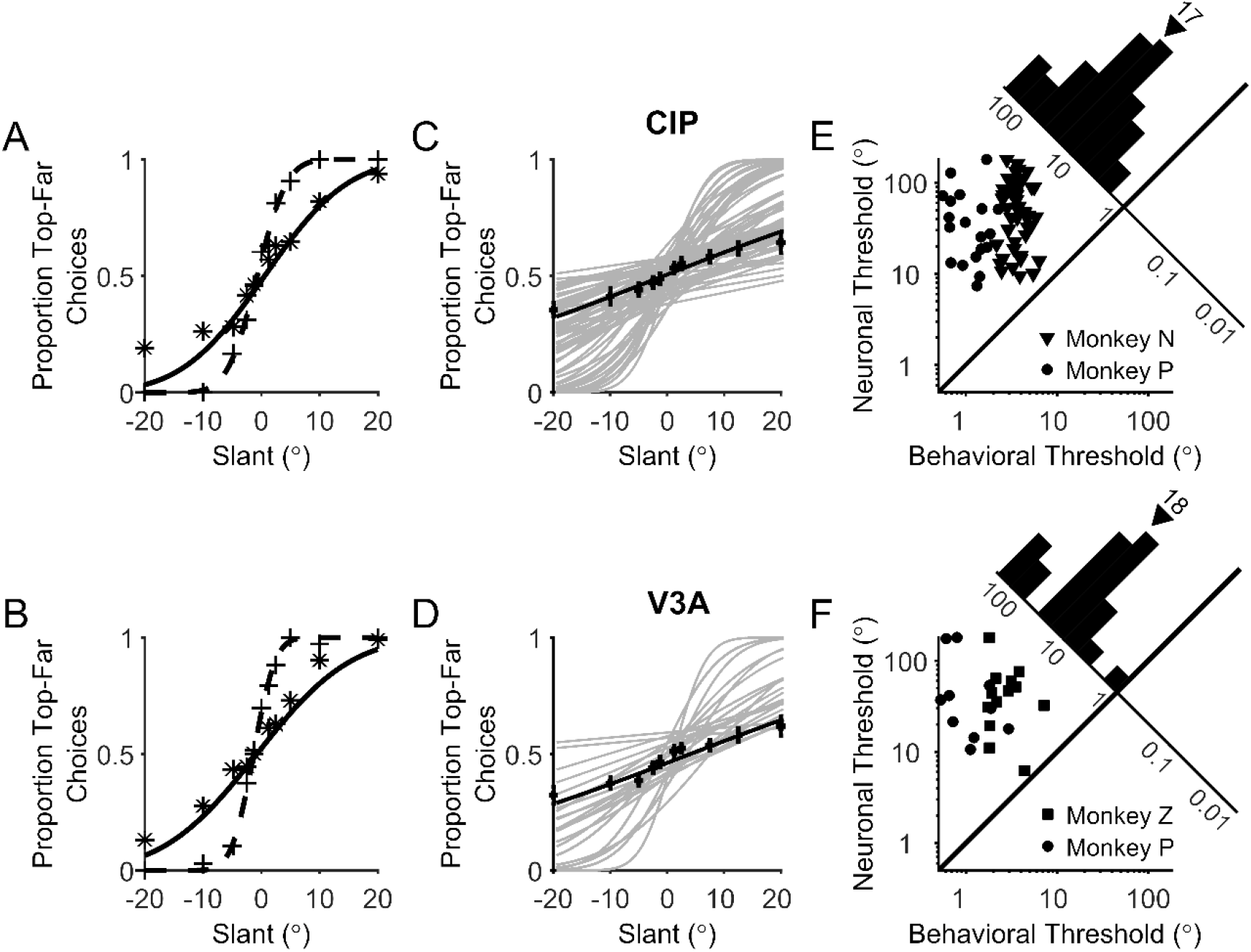
Comparison of behavioral and neuronal sensitivity. ***A & B***, The proportion of ‘top-far’ choices made during the recordings of the CIP (***A***) and V3A (***B***) neurons from Fig. 3 are plotted as a function of slant (+’s). Simultaneously measured neuronal responses were converted into neurometric functions using ROC analysis and the proportion of ‘top-far’ choices of an ideal observer are plotted as a function of slant (^*’^s). Dashed and solid curves show cumulative Gaussian fits to the psychometric and neurometric functions, respectively. ***C & D***, Gray curves show cumulative Gaussian fits to the neurometric functions of each neuron recorded during the slant discrimination task. Black symbols and curves show average neurometric functions across animals and neurons. Error bars denote SEM. ***E & F***, Behavioral and neuronal thresholds are compared for all individual experiments for monkeys N (triangles), P (circles), and Z (squares) for CIP (***E***) and V3A (***F***). Neuronal thresholds are multiplied by 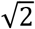 to account for the neuron/antineuron comparison. Diagonal histograms show distributions of neuronal to behavioral threshold ratios. Triangles above the histograms mark median threshold ratios.

To quantify the relationship between neuronal response and choice, choice probabilities (CPs) were computed using ROC analysis. For each slant, neuronal responses were grouped according to the choice. ‘Preferred’ choices corresponded to those made in favor of the neuron’s preferred slant, as determined from the 3D surface orientation tuning profile measured during fixation. ‘Non-preferred’ choices corresponded to those made in the opposite direction. CP was computed by performing ROC analysis on the preferred and non-preferred choice distributions for the (ambiguous) 0° slant stimulus. To achieve greater statistical power, a grand CP was computed by performing ROC analysis after normalizing the neuronal responses for each stimulus slant and combining the normalized data into two composite distributions corresponding to preferred versus non-preferred choices (Kang and Maunsell, 2012). Only stimulus slants for which the monkey made at least 3 choices in each direction were included in the grand CP calculation. To test if CPs were significantly different from chance level (CP = 0.50), a permutation test was used (1,000 permutations). The time course of choice-related activity was measured by computing CPs in 200 ms time windows shifted every 50 ms over the 1,000 ms stimulus duration.

To quantify the contributions of stimulus slant and choice to the responses of each neuron, Pearson correlations were computed between the following variables: slant, choice, and neuronal spike count. From these correlations, we computed a slant partial correlation, *r_FS.C_* (Eq. 2), that quantifies the relationship between spike count (F) and slant (S) while controlling for choice (C), and a choice partial correlation, *r_FC.S_* (Eq. 3), that quantifies the relationship between spike count and choice while controlling for slant. Because this analysis assumes a linear relationship between the stimulus and firing rate over the range of tested slants, we confirmed that the pattern of results did not change if slant was replaced with a nonlinear slant function including cubic, exponential, and sigmoidal functions, or if a larger partial correlation analysis was run which included multiple slant functions including the linear term. We did not consider nonlinear functions of choice because choice was a binary variable. Because the pattern of results did not depend appreciably on the stimulus function, as also reported recently for heading discrimination in the ventral intraparietal area (Zaidel et al., 2017), only the partial correlation analysis performed with slant, choice, and spike count is presented.

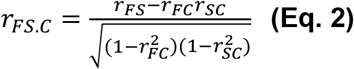

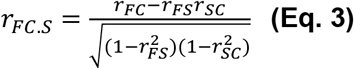

Positive slant partial correlations indicate that spike counts were greater for positive slants than negative slants. Positive choice partial correlations indicate that spike counts were greater for top-far than top-near choices. Partial correlations were computed based on spike counts over the entire 1,000 ms stimulus duration, as well as in 200 ms time windows shifted every 50 ms. For the partial correlation time course analysis, partial correlations were squared to determine how much variance in the spike counts was accounted for by the stimulus and the choice.

## Results

### Comparison of CIP and V3A responses to 3D surface orientation

Surface orientation tuning was measured for 427 CIP and 72 V3A neurons during a fixation task in which a checkerboard plane was presented at 25 slant–tilt combinations (Fig. 1A). Of these, 396 CIP (93%) and 60 V3A (83%) neurons were held for enough repetitions (≥ 3) to assess tuning. Tuning strength was quantified using a surface orientation discrimination index (SODI; see Materials and Methods) which ranges from 0 to 1. Larger SODI values indicate stronger tuning. The mean SODI in CIP was 0.63 ± 0.005 SEM (N = 396; Fig. 1B), and in V3A it was 0.68 ± 0.02 SEM (N = 60; Fig. 1C). The mean SODI was significantly smaller in CIP than V3A (Wilcoxon rank sum test, *p* = 5.8×10^−4^).

A two-step procedure was used to classify neurons as tuned or untuned. First, a one-way ANOVA was performed on the firing rates in response to each of the 25 slant–tilt combinations. Second, the tuning curve of each neuron that passed the ANOVA (p < 0.05) was fit with a Bingham function (Rosenberg et al., 2013). The second step eliminates neurons with multiple tuning peaks which would pass an ANOVA but are not selective for a unique stimulus (see Figure 5 in Rosenberg et al., 2013). Neurons with a Pearson correlation for the Bingham fit ≥ 0.8 were classified as tuned, and otherwise untuned. Based on these criteria, 215 CIP neurons (54% of the 396 tested) and 44 V3A neurons (73% of the 60 tested) were tuned. Of the neurons classified as untuned, 26.5% in CIP (48/181) and 25% in V3A (4/16) were rejected for having multiple peaks.

**Figure 5.**
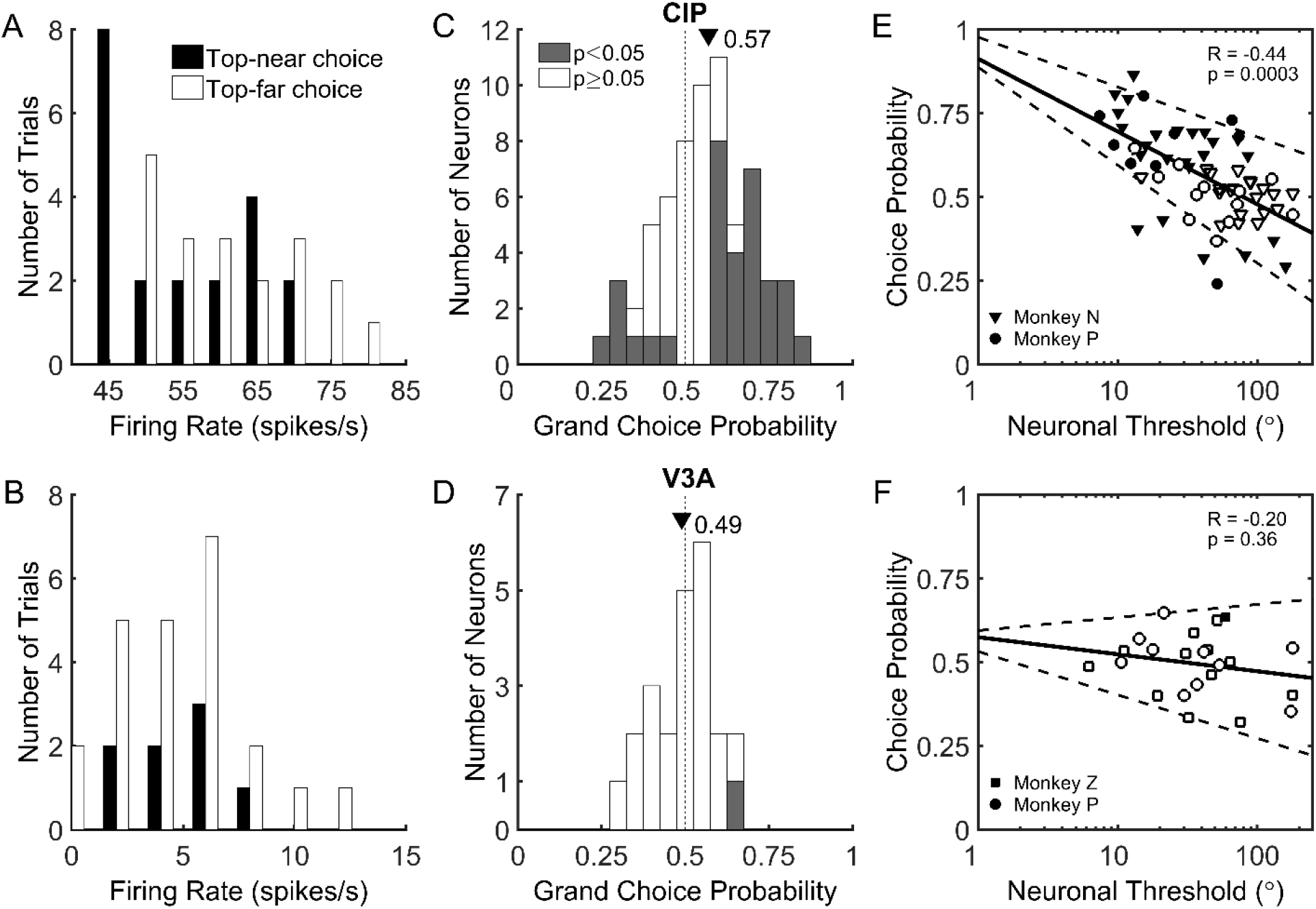
Summary of choice-related activity in CIP and V3A. ***A & B***, Distribution of firing rates for example CIP (***A***) and V3A (***B***) neurons (same as in Figs. 3, 4A,B) in response to the ambiguous 0° slant stimulus. Responses are sorted according to whether the monkey made a ‘top-near’ (black) or ‘top-far’ (white) choice. For the CIP neuron, the choice-related difference in responses yielded a choice probability significantly different from chance (grand C*p* < 0.65, *p* = 0.001). For the V3A neuron, there was no significant choice-related difference in responses (grand C*p* = 0.50, *p* = 0.36). ***C & D***, Histograms of grand choice probabilities for all 65 CIP (***C***) and 23 V3A (***D***) neurons. Gray bars denote CPs that are significantly different from the chance value of 0.50 (p < 0.05, permutation test). Mean CPs are marked by triangles. ***E & F***, Choice probability as a function of neuronal threshold (multiplied by 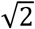). There is a significant negative correlation between CP and neuronal threshold in CIP (***E***) and no significant correlation between CP and neuronal threshold in V3A (***F***). Solid lines show linear fits and dashed lines show 95% confidence intervals for the slope. Filled symbols denote CPs significantly different from chance (0.50, *p* < 0.05, permutation test). Different symbols correspond to different animals.

The distribution of slant–tilt preferences was examined for each area by performing an equal area preserving projection (Rosenberg et al., 2013) and plotting the preferred slant and tilt of each neuron in that space (Fig. 1D). We previously found that the distribution of CIP slant–tilt preferences was not significantly different from uniform in untrained animals (Rosenberg et al., 2013). Here we found that the distribution of preferences in CIP and V3A were significantly different from uniform (Chi-squared test, CIP: *p* = 1.07×10^−7^; V3A: *p* = 0.01). In particular, there was a bias towards representing smaller slants (note the relative sparsity of cells near the top of the scatter plot in Fig. 1D). It is possible that extensive training in the fine slant discrimination task resulted in a shift in tuning preferences towards smaller slants.

### Slant discrimination behavior

A control experiment was conducted to confirm that the animals did not perform the slant discrimination task based on local absolute disparity cues signaling that the upper (lower) half of the plane was in front of (behind) the LCD. Each monkey performed the slant discrimination task for nine sessions with the stimuli centered at three depths (0 and ±2.25 cm) from the display (Fig. 2A,B). Psychometric functions for each monkey and depth are shown in Fig. 2C. The proportion of ‘top-far’ choices is plotted for each slant and fit with a cumulative Gaussian function. One-way ANOVAs showed no significant effect of depth on the point of subjective equality (P.S.E.; monkey N: *F* = 0.65, *p* = 0.53; monkey P: *F* = 0.53, *p* = 0.60; monkey Z: *F* = 2.41, *p* = 0.12) or threshold (monkey N: *F* = 0.58, *p* = 0.57; monkey P: *F* = 0.11, *p* = 0.90; monkey Z: *F* = 0.70, *p* = 0.51). Although not significant, there was a slight tendency for the P.S.E. to be negative at −2.25 cm (Fig. 2D). However, if the animals were relying on local absolute disparity cues to perform the task, the P.S.E. would have a magnitude of at least 14° at the near/far depths (i.e., the smallest slant at which a plane would cross the screen), which is much greater than the average P.S.E. of −0.38° at −2.25 cm. These data strongly suggest that the monkeys performed the task by assessing the slant of the plane rather than by judging local stimulus depth relative to the plane of fixation.

### Neuronal sensitivity during slant discrimination

Of the 215 tuned CIP neurons, 151 (70%) were significantly tuned for slant (ANOVA, *p* < 0.05) along the 90°/270° tilt axis used in the slant discrimination task (white dashed lines in Fig. 3A,B), and therefore studied further. Of these, data from 65 (43%) were included in this study. The remaining 86 neurons (57%) were recorded for another task (16 neurons, 11%) or were not recorded for a sufficient number of repetitions (≥ 10) to be included (70 neurons, 46%). Likewise, of the 44 V3A neurons, 35 (80%) were significantly tuned for slant along the 90°/270° tilt axis. Of these, 23 (66%) were held for sufficient repetitions (≥ 10) to be included.

Surface orientation tuning curves for example CIP and V3A neurons that met these criteria are shown in Fig. 3A,B. Responses recorded during the slant discrimination task are shown in Fig. 3C,D for the same neurons. For both neurons, tuning was monotonic over the range of slants presented in the discrimination task. The CIP neuron (Fig. 3C) fired more in response to positive slants (top of the plane further from the animal), whereas the V3A neuron (Fig. 3D) fired more in response to negative slants (top of the plane closer to the animal).

To assess how well the responses of these neurons could be used to discriminate slants of opposite sign, we compared firing rate distributions. Firing rate distributions for 3 pairs of slants (±20°, ±5°, and ±1.25°) are shown in Fig. 3E,F. Note that the distributions are completely overlapping for ±1.25° whereas there is little or no overlap at ±20°. Consequently, an ideal observer could reliably discriminate ±20° of slant based on the firing rates of these neurons, but would be unable to discriminate ±1.25° of slant. The ability of an ideal observer to discriminate slants of opposite sign was quantified using ROC analysis (Britten et al., 1996; Gu et al., 2007). The probability that an ideal observer could correctly report whether the slant of a presented plane was positive or negative was calculated for each slant magnitude. A neurometric function was then constructed by plotting ROC values for each slant pair (i.e., positive and negative slants of the same magnitude), and fitting the function with a cumulative Gaussian (Fig. 4A,B, solid curves). A neuronal threshold quantifying the neuron’s sensitivity to changes in slant was defined as the standard deviation of the cumulative Gaussian fit. This analysis was performed for each of the 65 CIP and 23 V3A neurons, and the resulting neurometric functions are shown in Fig. 4C,D. Across all monkeys, median neuronal thresholds were 32.86° in CIP and 26.25° in V3A, and were not significantly different (Wilcoxon rank sum test, *p* = 0.48). We further confirmed that neuronal thresholds were similar between monkeys. The median CIP thresholds were 35.16° (monkey N) and 26.04° (monkey P), and not significantly different (Wilcoxon rank sum test, *p* = 0.30). Likewise, the median V3A thresholds were 31.30° (monkey Z) and 23.73° (monkey P), and not significantly different (Wilcoxon rank sum test, *p* = 0.58). These results indicate that CIP and V3A neurons are similarly sensitive to changes in slant.

Neurometric functions can be directly compared to psychometric functions measured in the same recording session (dashed curves, Fig. 4A,B). Simultaneously measured neuronal and behavioral thresholds are compared in Fig. 4E,F for CIP and V3A, respectively. For this comparison, neurometric thresholds were multiplied by 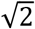 since the neurometric functions were constructed by comparing two response distributions (the neuron/anti-neuron approach), whereas the behavioral task had a single stimulus interval. Distributions of neuronal to behavioral threshold ratios are shown as diagonal histograms. All of the neuronal/behavioral threshold ratios were greater than 1, indicating that no CIP or V3A neuron was more sensitive than a monkey. Monkey N’s median neuronal/behavioral threshold ratio was 14 for CIP, monkey P’s median threshold ratio was 34 for CIP and 30 for V3A, and monkey Z’s median threshold ratio was 16 for V3A. Although behavioral sensitivity was generally greater than neuronal sensitivity, the thresholds of some neurons approached that of the behavior, suggesting that CIP and V3A could contribute to performance of the slant discrimination task.

### Neuronal responses in CIP but not V3A correlate with slant reports

During the slant discrimination task, variability was observed in both the neuronal firing rates and choices elicited by stimuli of the same slant. This variability is evident in histograms of the example CIP neuron’s responses to a slant of 0°, grouped by choice (Fig. 5A). This stimulus is ambiguous and there is no correct answer because the top of the plane leans neither toward nor away from the monkey. Thus, the monkey made choices toward both response targets with roughly equal frequency. For the example CIP neuron, the firing rate tended to be lower when the monkey made a top-near choice and greater when the monkey made a top-far choice. In other words, responses were greater when the monkey chose the target corresponding to the neuron’s slant preference. In contrast, the example V3A neuron preferred negative slants, but the histograms of responses to a slant of 0°, grouped by choice, were largely overlapping. Thus, there was no clear difference in the activity of the example V3A neuron when the animal made top-far versus top-near choices (Fig. 5B).

Choice probability (CP) analysis was used to quantify the relationship between neuronal response and choice (Celebrini and Newsome, 1994; Britten et al., 1996; Dodd et al., 2001; Nienborg and Cumming, 2006; Gu et al., 2007). We computed the CP by first assigning neuronal responses to two groups according to the animal’s choice. ‘Preferred’ slant choices were made in the direction of the preferred slant and ‘non-preferred’ slant choices were made in the direction of the non-preferred slant. Preferred and non-preferred slants were defined according to the tuning preference along the 90°/270° tilt axis that was measured during the 3D orientation tuning (fixation only) task. Slant preferences generally matched between the fixation and discrimination tasks, with the preference reversing for only 6 CIP neurons and 1 V3A neuron. Since reversals of slant preference could be an effect of choice-related signals during the discrimination task, we computed CPs based on stimulus preferences measured during fixation.

After sorting responses by choice, we used ROC analysis to compute the probability that an ideal observer could predict the monkey’s choice based on the neuron’s responses (see Materials and Methods). The CP was calculated in two ways. First, we only considered responses to the ambiguous 0° slant stimulus. For the CIP neuron in Fig. 5A, the CP was 0.65, indicating it fired more when the monkey made a choice in favor of the preferred slant. Across all CIP neurons, the mean CP for a 0° slant stimulus was 0.58, which was significantly greater than the chance value of 0.50 (*t*-test, *t* = 3.89, *p* = 2.45×10^−4^). For the V3A neuron in Fig. 5B, the CP was 0.45, suggesting the neuron fired slightly more when the monkey made a choice in favor of the cell’s non-preferred slant. Across all V3A neurons, the mean CP for the 0° slant stimulus was 0.52, which was not significantly different from chance (*t*-test, *t* = 0.64, *p* = 0.53). Second, to achieve greater statistical power, we calculated a ‘grand CP’ by including responses to all slants for which the monkey made at least 3 choices toward each response target. For this analysis, responses to each slant were normalized using the balanced Z-score method (Kang and Maunsell, 2012). For the CIP neuron in Fig. 5A, the grand CP was 0.65 and significantly greater than the chance value of 0.50 (permutation test, 1000 permutations, *p* = 0.001). The grand CP for the V3A neuron in Fig. 5B was 0.50 and not significantly different from chance (p = 0.36). Across the neural populations, the grand CP was highly correlated with the CP measured for the 0° slant stimulus (CIP: *r* = 0.81, *p* = 1.0×10^−15^; V3A: *r =* 0.78, *p* = 0.0001). The analyses that follow are based on grand CPs.

Fig. 5C,D shows CP histograms for CIP and V3A. The mean CIP CP was 0.57, which was significantly greater than 0.50 (*t*-test, *p* = 1×10^−15^). The mean CIP CP was also significantly different from chance for each monkey (*t*-test, monkey N: CP = 0.57, *p* = 3.40×10^−4^; monkey P: CP = 0.57, *p* = 0.04). In total, 51% of CIP neurons (33/65) had CPs that were significantly different from chance (permutation test, 1000 permutations, *p* < 0.05). For the majority of CIP neurons with significant CPs (26/33), firing rates increased when the monkey made a choice in favor of the preferred slant (CPs > 0.50). However, 7 CIP CPs were significantly below 0.50, indicating they fired more when the monkey made a choice in favor of the non-preferred slant. In contrast to CIP, the mean V3A CP was 0.48, which was not significantly different from 0.50 (*t*-test, *p* = 0.40). Neither monkey had a mean V3A CP that was significantly different from chance (*t*-test, monkey P: CP = 0.48, *p* = 0.42; monkey Z: CP = 0.49, *p* = 0.67). Permutation tests revealed that only one V3A neuron had a CP that was significantly different from chance. As a control, we confirmed that there was no significant difference in CP associated with whether the neurons preferred positive or negative slants. The mean CIP CP was 0.55 ± 0.03 SEM (N = 30) for neurons preferring positive slants and 0.59 ± 0.03 SEM (N = 35) for those preferring negative slants (*t*-test, *t* = 1.49, *p* = 0.14). The mean V3A CP was 0.46 ± 0.03 SEM (N = 10) for neurons preferring positive slants and 0.50 ± 0.03 SEM (N = 13) for those preferring negative slants (*t*-test, *t* = 1.29, *p* = 0.21). Comparing choice-related activity across the two areas, we found that the mean CIP CP was significantly greater than the mean V3A CP (*t*-test, *p* = 0.003). These findings indicate that CIP, but not V3A, neurons display strong choice related activity during the slant discrimination task.

We further found that CIP neurons showed a significant negative correlation between neuronal threshold and CP (*r* = −0.44, *p* = 3×10^−4^; Fig. 5E). The 10 most sensitive CIP neurons had a mean CP of 0.72 ± 0.03 (SEM), whereas the 10 least sensitive had a mean CP of 0.48 ± 0.03 (SEM). In contrast, the correlation between neuronal threshold and CP was not significant in V3A (*r* = −0.20, *p* = 0.36), and the V3A CPs clustered around 0.50 regardless of neuronal threshold (Fig. 5F). We additionally ran an ANCOVA in which CP was the dependent variable, neuronal threshold was a continuous covariate, and the brain area was an ordinal factor. We found a significant interaction (*p* = 0.03) between neuronal threshold and brain area, indicating a significant difference in the strength of the relationship between CP and neuronal threshold in CIP and V3A.

As a control, we confirmed that trial-by-trial variation in vertical eye position, vertical eye velocity, and vergence during the stimulus presentation had no appreciable effect on CIP CPs and neuronal thresholds (Gu et al., 2007). For each CIP neuron, we performed three separate analyses of covariance (ANCOVAs) to test the relationship between neuronal firing rate and choice with vertical eye position, vertical eye velocity, or vergence as co-regressors (averaged over the length of each trial). Fifteen percent (10/65) of CIP neurons had a significant dependence of firing rate on vertical eye position, 3% (2/65) had a significant dependence of firing rate on vertical eye velocity, and 6% (4/65) had a significant dependence of firing rate on vergence (p < 0.05, ANCOVA, Bonferroni-Holm correction for multiple comparisons). We therefore calculated CPs and neuronal thresholds after removing the dependence (linear trend) on vertical eye position, vertical eye velocity, and vergence from the neuronal responses. After removing the effect of vertical eye position, there was a small but significant reduction in CP (0.57 before versus 0.56 after correction; paired *t*-test, *t* = 2.53, *p* = 0.01). The CP measurements before and after correction were highly correlated (*r* = 0.96, *p* = 1.0×10^−16^), and the mean value remained significantly greater than chance after correction (*t*-test, *t* = 3.61, *p* = 5.95×10^−4^). Removal of the effect of vertical eye position had no significant effect on the median neuronal threshold (Wilcoxon sign-rank test, *p* = 0.24). For vertical eye velocity, there was a small but significant effect on the mean CP (0.57 before versus 0.56 after correction, paired *t*-test, *t* = 3.05, *p* = 0.003) and the median neuronal threshold (32.86° before versus 38.06° after correction, Wilcoxon sign-rank test, *p* = 0.03). The CP measurements before and after correction were highly correlated (*r* = 0.95, *p* = 1.0×10^−16^), and remained significantly greater than chance after correction (*t*-test, *t* = 3.57, *p* = 6.92×10^−4^). Neuronal thresholds were also highly correlated before and after correction (*r* = 0.85, *p* = 3.0×10^−15^). For vergence, there was no significant effect on mean CP (*p* = 0.58) or median neuronal threshold (*p* = 0.48). Thus, variations in eye position, eye velocity, and vergence had little effect on CIP CPs and neuronal thresholds.

### Contributions of stimulus and choice to CIP and V3A responses

During the slant discrimination task, both the stimulus and the choice may contribute to neuronal activity. The contributions of stimulus and choice to the activity of example CIP and V3A neurons is shown in Fig. 6. Slant tuning curves measured by averaging firing rates across all presentations of each slant without regard to the choice are shown in black. For comparison, choice-conditioned slant tuning curves were computed for top-far and top-near choices (orange and purple curves in Fig. 6, respectively). Only slants for which the monkey made at least three choices in the relevant direction were included in the choice-conditioned tuning curves. In CIP, choice-conditioned tuning curves often showed clear separation, indicating a strong effect of choice on firing rate. For the CIP neuron in Fig. 6A, the top-far choice-conditioned tuning curve (orange) lies above the top-near choice-conditioned tuning curve (purple). This difference indicates the neuron responded more strongly when the monkey made a choice in the direction of the neuron’s preferred slant (top-far). Correspondingly, the neuron’s CP is greater than 0.50. In contrast, Fig. 6B shows a CIP neuron that responded more strongly when the monkey made a choice in the opposite direction of the preferred slant. Hence, the top-near choice-conditioned tuning curve (purple) is above the top-far choice-conditioned tuning curve (orange), and the CP is less than 0.50. In V3A, choice-conditioned tuning curves largely overlapped. This was the case even when the CP was relatively large, as shown for the neuron in Fig. 6C, indicating that choice had little effect on V3A responses.

**Figure 6.**
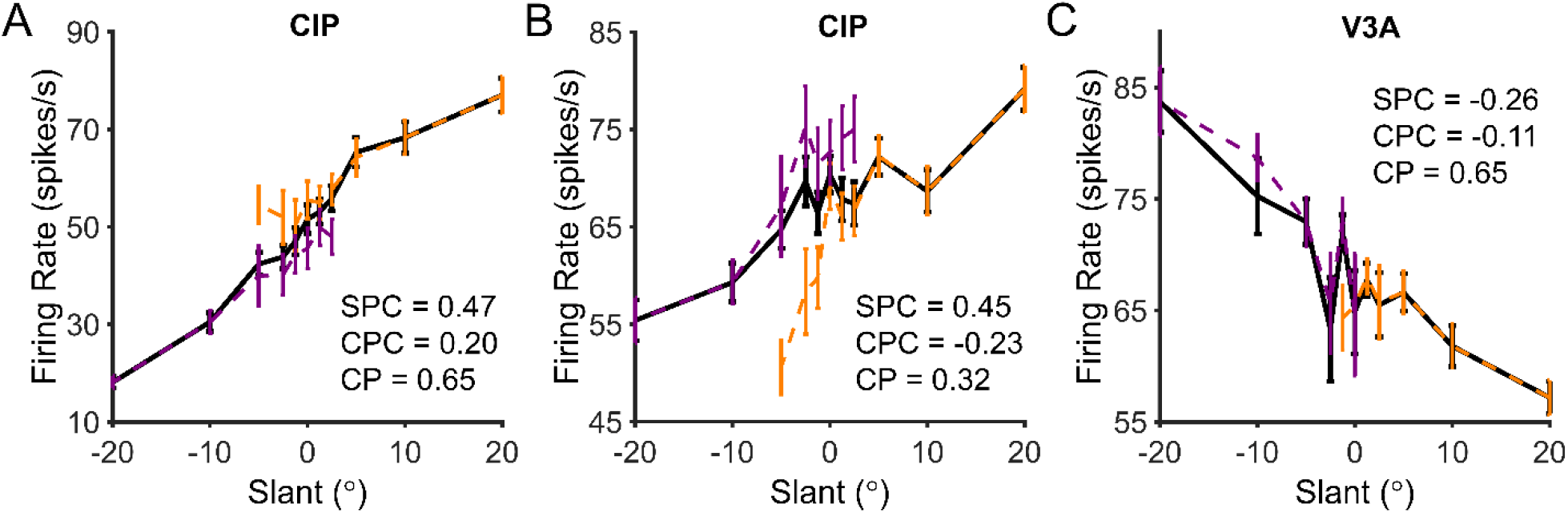
Example CIP and V3A neurons illustrating the effect of choice on slant tuning. For each neuron, the black curve shows the slant tuning curve created by averaging responses regardless of choice. Orange and purple curves show choice-conditioned slant tuning curves created by separating responses into top-far versus top-near choices, respectively. The slant partial correlation (SPC), choice partial correlation (CPC), and choice probability (CP) are listed for each neuron. ***A***, CIP neuron with a positive SPC, a positive CPC, and a CP > 0.50 (*p* = 0.001). ***B***, CIP neuron with a positive SPC, a negative CPC, and a CP < 0.50 (*p* = 0.001). ***C***, V3A neuron with a negative SPC, a negative CPC, and a CP > 0.50 (*p* = 0.29).

To dissociate the contributions of stimulus and choice to each neuron’s responses, partial correlations were computed between slant, choice, and spike counts using all trials. This analysis estimates how much variance in the responses can be accounted for by stimulus and choice while controlling for the fact that these variables are correlated. Similar percentages of CIP (30/65; 46%) and V3A (10/23; 43%) neurons had significant slant partial correlations (*p* < 0.05), and the magnitude (absolute value) of the slant partial correlations in CIP (median = 0.09) and V3A (median = 0.15) were not significantly different (Wilcoxon rank-sum test, *p =* 0.14). The ranges of slant partial correlations in CIP (*r* = −0.51 to 0.47) and V3A (*r* = −0.48 to 0.46) were also similar. Correspondingly, the variance of the slant partial correlations was not significantly different between the areas (Levene’s test, *W* = 2.11, *p* = 0.15).

Although the slant partial correlations in CIP and V3A were similar, the choice partial correlations differed substantially. A greater percentage of neurons had significant choice partial correlations in CIP (40/65; 62%) than V3A (7/23; 30%), and the magnitude of the choice partial correlations in CIP (median = 0.13) was significantly greater than in V3A (median = 0.09), Wilcoxon rank-sum test (*p* = 0.003). The range of choice partial correlations was also greater in CIP (*r* = −0.55 to 0.49) than V3A (*r* = −0.15 to 0.20). Correspondingly, the variance of the choice partial correlations was significantly different between the areas (Levene’s test, *W* = 9.19, *p* = 0.003). These findings confirm that choice had a greater effect on CIP than V3A activity.

In CIP, the relative signs of the slant and choice partial correlations were largely predictive of CP. The CIP neuron in Fig. 6A preferred positive slants (positive slant partial correlation) and top-far choices (positive choice partial correlation). Consistent with this, the CP was significantly greater than 0.50 (*p* = 0.001). In contrast, the CIP neuron in Fig. 6B preferred positive slants (positive slant partial correlation) but top-near choices (negative choice partial correlation). Consistent with this, the CP was significantly less than 0.50 (*p* = 0.001). For comparison, a V3A neuron that preferred negative slants and top-near choices is shown in Fig. 6C. Although the CP was greater than 0.50, it was not significantly different from 0.50 (*p* = 0.29).

The relationships between slant partial correlation, choice partial correlation, and CP are summarized for CIP and V3A in Fig. 7. Quadrant I (upper right) contains neurons for which positive slants and top-far choices increased firing rate. Quadrant III (lower left) contains neurons for which negative slants and top-near choices increased firing rate. Note that top-far (top-near) choices were correct for positive (negative) slants; thus, quadrants I and III contain neurons with congruent stimulus and choice effects. Based on the example cells in Fig. 6, quadrants I and III should contain neurons with CPs greater than 0.50, at least in CIP where choice effects are robust. Consistent with this prediction, the CP of 35/40 (88%) of the CIP neurons in quadrants I and III was greater than 0.50 (Fig. 7A) and the mean CP was 0.59 ± 0.02 (SEM, N = 40), which was significantly greater than 0.50 (Wilcoxon signed rank test, *p* = 3.3×10^−6^).

**Figure 7.**
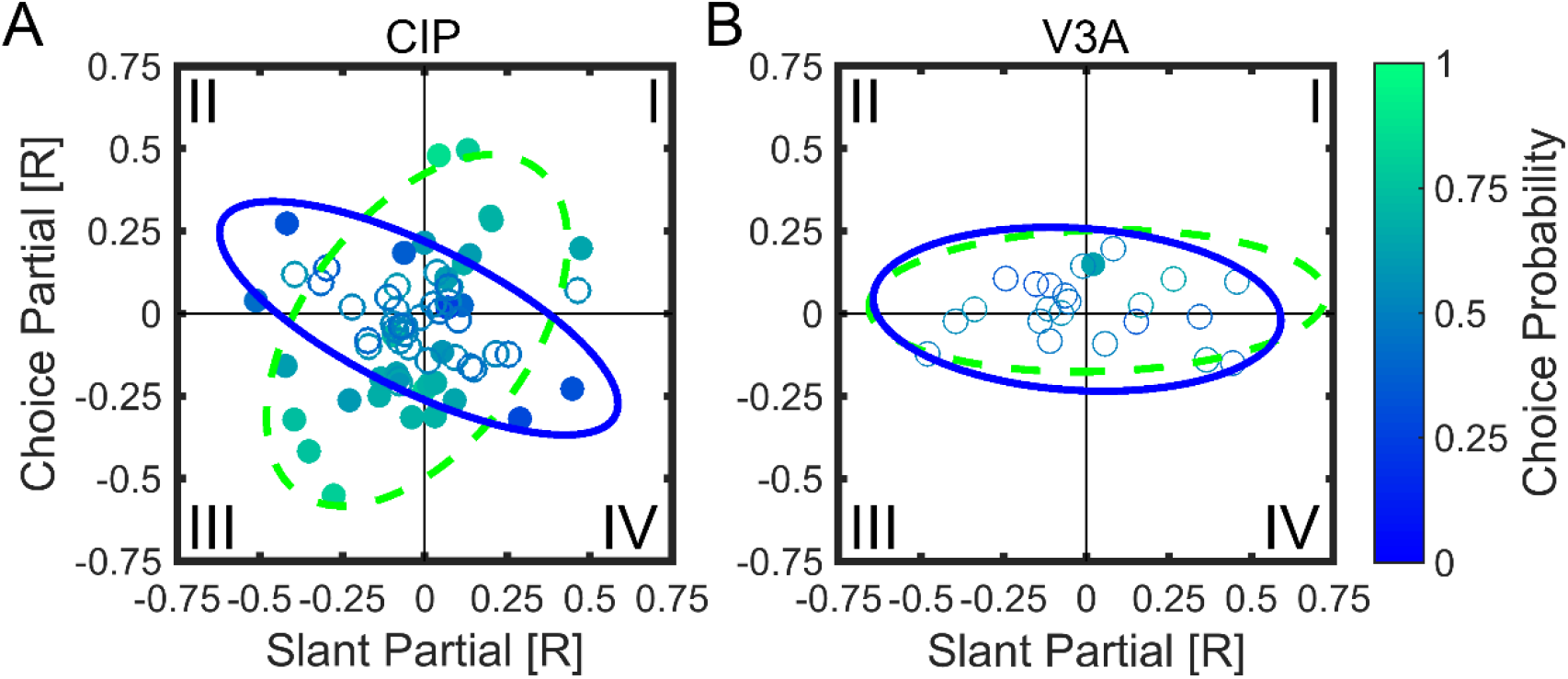
Partial correlation analysis showing relationships between slant partial correlation, choice partial correlation, and CP. Choice partial correlation is plotted as a function of slant partial correlation with individual neurons color coded to indicate CP. Significant CPs are filled, nonsignificant CPs are open. Data are shown for 65 CIP (***A***) and 23 V3A (***B***) neurons. Curves show 95% confidence ellipses fit to data points with CP > 0.50 (green dashed) or CP < 0.50 (blue solid). ***A***, In CIP, as indicated by the oblique orientations of the 95% confidence ellipses, CPs > 0.50 (greener) tended to occur when the slant and choice partial correlations had the same sign (quadrants I and III) whereas CPs < 0.50 (bluer) tended to occur when the slant and choice partial correlations had opposite signs (quadrants II and IV). ***B***, For V3A, choice-related activity was weak, as indicated by the elongated but horizontally oriented 95% confidence ellipses.

There was also a substantial number of neurons for which slant and choice had opposite effects on firing rate (quadrants II and IV). Cells in quadrant II (upper left) are those for which firing rate increased for negative slants and top-far choices. Cells in quadrant IV (lower right) are those for which firing rate increased for positive slants and top-near choices. Assuming that CP was computed based on the true sign of the slant preference (determined from the surface orientation tuning curve measured during fixation to minimize choice-related activity; the sign reversed for one CIP neuron in quadrants II/IV if determined from the slant discrimination data), neurons in quadrants II and IV should have CPs less than 0.50. This was not immediately evident: 12/25 (48%) CIP neurons in these quadrants had CPs less than 0.50, and the mean Cp = 0.50 ± 0.02 SEM was not significantly different from 0.50 (N = 25, Wilcoxon signed rank test, *p* = 0.95). Note, however, that neurons with the lowest CPs (darker blue points) are largely found in quadrants II and IV.

Thus, to further test if CPs are related to the relative signs of the slant and choice partial correlations, we fit a 95% confidence ellipse to the data from all CIP neurons with CPs > 0.50 (green dashed ellipse) and a 95% confidence ellipse to those with CPs < 0.50 (blue solid ellipse), as shown in Fig. 7A. Consistent with our predictions, the ellipses are obliquely oriented and nearly orthogonal. The orientation of the major axis for the CPs > 0.50 ellipse is 53.71° with a bootstrapped 95% confidence interval of [34.92° 68.60°], indicating it is elongated along quadrants I and III. The orientation of the major axis for the CPs < 0.50 ellipse is 153.67° with a bootstrapped 95% confidence interval of [142.53° 165.04°], indicating it is elongated along quadrants II and IV. Thus, in CIP, neurons with CPs > 0.50 tend to have slant and choice partial correlations of the same sign, whereas neurons with CPs < 0.50 tend to have slant and choice partial correlations of opposite sign. The slant and choice partial correlations in CIP were not significantly correlated with each other overall (*r* = −0.17, *p* = 0.18), suggesting that slant and choice can have independent effects on neuronal responses (see Discussion).

In V3A, the mean CP for quadrants I and III (0.55 ± 0.03 SEM) was not significantly greater than 0.50 (N = 9, Wilcoxon signed rank test, *p* = 0.09), but the mean CP for quadrants II and IV (0.44 ± 0.02 SEM) was significantly less than 0.50 (N = 14, Wilcoxon signed rank test, *p* = 0.02). This suggests there was some tendency for the relative signs of the slant and choice partial correlations to predict CP in V3A. However, this trend was weak compared to CIP, as demonstrated by the 95% confidence ellipses for CPs > 0.50 and CPs < 0.50 in V3A. For both ellipses, the major axis is oriented approximately along the slant partial correlation axis (1.08° and −3.53° for CPs > 0.50 and CPs < 0.50, respectively), reflecting that V3A responses were substantially more dependent on slant than choice.

### Time course of stimulus-related and choice-related activity in CIP and V3A

Lastly, we examined the time course of CPs, neuronal thresholds, and partial correlations in CIP and V3A by computing these quantities within a series of 200 ms bins shifted every 50 ms. Average CP time courses are shown in Fig. 8A,B for CIP and V3A, respectively. The mean CIP CP increased above baseline relatively late in the stimulus duration and remained elevated. The first time bin in which the mean CP (0.53 ± 0.01 SEM, N = 65 neurons) was significantly greater than 0.50 was 250 ms after stimulus onset (one-way ANOVA with multiple comparisons for N = 17 time bins, *p* < 0.05). The mean V3A CP was not significantly different from 0.50 in any time bin (one-way ANOVA with multiple comparisons for N = 17 time bins, *p* ≥ 0.05), but was slightly less than 0.50 throughout most of the stimulus duration. For comparison, mean CIP and V3A neuronal thresholds are shown in Fig. 8C,D, respectively.

**Figure 8.**
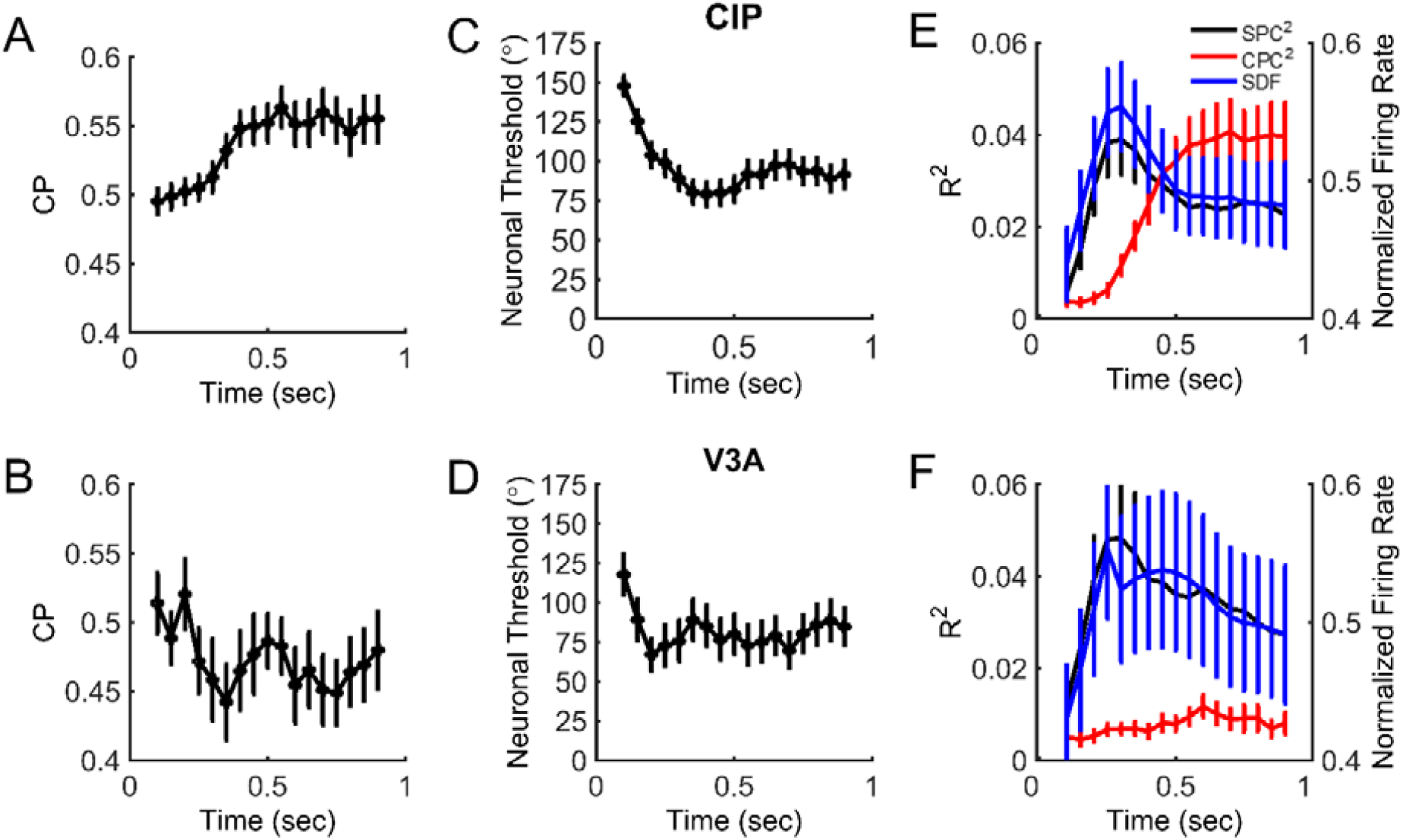
Time courses of choice probability, neuronal threshold, and partial correlations. ***A & B***, Mean values of choice probability (CP) for CIP (***A***) and V3A (***B***) neurons as a function of time relative to stimulus onset. ***C & D***, Mean neuronal thresholds (multiplied by 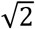) for CIP (***C***) and V3A (***D***) as a function of time. ***E & F***, Mean spike density functions (SDF, blue) as well as squared slant (SPC, black) and choice (CPC, red) partial correlations for CIP (***E***) and V3A (***F***) as a function of time. In all plots, analysis bins are 200 ms in duration, shifted every 50 ms starting at 100 ms. Each point is plotted in the center of the 200 ms time bin. Error bars denote SEM.

The mean time courses for the spike density function (SDF; a measure of the average population response), squared slant partial correlation (SPC), and squared choice partial correlation (CPC) are shown for CIP and V3A in Fig. 8E,F, respectively. The time courses of the squared slant partial correlations (black curves) are highly similar to the mean spike density functions (blue curves), with an early peak and smaller sustained values. In fact, the time course of the spike density function was highly correlated with that of the slant partial correlation in both areas (CIP: *r* = 0.95, *p* = 5.2×10^−9^, N = 17; V3A: *r* = 0.90, *p* = 7.0×10^−7^, N = 17). In CIP, the time course of the squared slant partial correlation peaked around 150-200 ms, whereas the squared choice partial correlation increased later in the stimulus duration (red curve). It was not until 350 ms after stimulus onset that the squared choice partial correlation became significantly different from its initial value (one-way ANOVA with multiple comparisons, *p* < 0.05), further emphasizing that choice-related activity in CIP is substantially delayed relative to stimulus-related activity. However, for V3A, the squared choice partial correlation remained close to zero throughout the stimulus duration and showed no significant difference from the initial value at any time point (oneway ANOVA with multiple comparison, *p* = 0.42), further reflecting that there was little to no choice-related activity in V3A during the slant discrimination task.

## DISCUSSION

We investigated correlations between 3D surface orientation perception and neuronal activity in areas V3A and CIP of the macaque monkey. Our results show that surface orientation is similarly discriminable based on V3A and CIP responses, and that neurons in the two areas are similarly sensitive to small slant variations. Together with anatomical data (Nakamura et al., 2001), these results suggest V3A may, at least partially, underlie 3D orientation selectivity in CIP (Taira et al., 2000; Tsutsui et al., 2001; Rosenberg et al., 2013; Rosenberg and Angelaki, 2014b). Although stimulus-related activity was similar in the two areas, choice-related activity differed qualitatively. Specifically, choice-related activity during the slant discrimination task was prominent in CIP but largely lacking in V3A, implying a functional distinction between the areas. Together, these results suggest that both areas may contribute to 3D surface orientation processing, but only CIP carries 3D orientation decision-related signals.

### Comparison of stimulus-related and choice-related activity in CIP and V3A

The present results strongly agree with previous reports of 3D orientation selectivity in CIP (Taira et al., 2000; Rosenberg et al., 2013), and are consistent with previous studies implicating V3A in binocular disparity processing, 3D vision, and prehensile sensorimotor processing (Nakamura et al., 2001; Tsao et al., 2003; Anzai et al., 2011; Ban and Welchman, 2015; Goncalves et al., 2015). In both areas, 3D orientation preferences were shifted towards small slant preferences. This non-uniformity differs from our previous finding of a uniform distribution of 3D orientation preferences in CIP (Rosenberg et al., 2013), and may be a byproduct of extensive slant discrimination training about the frontoparallel plane. Based on 3D orientation tuning measured during passive fixation, the strength of selectivity was similar between the two areas (quantified using the SODI), though slightly greater in V3A than CIP. When slant tuning was measured during the slant discrimination task, V3A and CIP neurons were similarly sensitive to small slant changes, as evidenced by similar average neuronal thresholds. For some neurons in each area, neuronal thresholds were nearly as small as the behavioral threshold, suggesting that the animals may be less sensitive to changes in slant than is possible from an optimal decoding of the neuronal activity. Recent theoretical work suggests that sub-optimal decoding and/or information-limiting noise correlations that introduce redundancy may cause behavioral thresholds to be only slightly smaller than individual neuronal thresholds (Moreno-Bote et al., 2014; Pitkow et al., 2015).

Although we found similar stimulus response properties in V3A and CIP, there was a stark difference in their choice-related activity. More than half of the CIP neurons had significant CPs, whereas only one V3A neuron had a significant CP. This difference indicates that CIP activity is functionally coupled with perceptual slant decisions, whereas V3A activity is not. However, the relationship between CIP activity and choice is not necessarily causal. Significant CPs could arise in a bottom-up manner (Britten et al., 1996; Haefner et al., 2013; Wimmer et al., 2015), but there is growing evidence that top-down (feedback) signals make important contributions to the presence of CPs (Nienborg and Cumming, 2009; Wimmer et al., 2015; Cumming and Nienborg, 2016; Kwon et al., 2016; Yang et al., 2016). Thus, observing significant choice-related activity does not necessarily imply a contribution to perceptual decisions, as reinforced by recent findings of dissociations between choice-related activity and reversible inactivation of brain areas. For example, macaque VIP neurons have substantially greater CPs than MSTd neurons during a heading discrimination task (Gu et al., 2007; Gu et al., 2008; Chen et al., 2013), yet inactivation of MSTd impairs task performance whereas inactivation of VIP does not (Gu et al., 2012; Chen et al., 2016). Similarly, macaque LIP shows robust choice-related activity during motion discrimination, but inactivation of LIP does not impair task performance (Katz et al., 2016). A causal relationship between 3D surface orientation perception and CIP activity thus remains uncertain.

Previous work has shown that the 3D orientation tuning of CIP neurons is largely invariant to changes in the mean depth of the stimuli relative to the fixation plane, as well as the defining visual (i.e., perspective or stereoscopic) cue, suggesting that CIP neurons are sensitive to depth gradients (Taira et al., 2000; Tsutsui et al., 2001; Rosenberg and Angelaki, 2014b). In the present study, we did not have sufficient stimulus conditions to determine whether the slant selectivity of V3A neurons is also robust to changes in mean depth. Thus, we cannot rule out the possibility that some of the selectivity we observed in V3A reflects local disparity selectivity, given that local disparity within the receptive field changes as a function of slant in our stimulus. Indeed, an intriguing hypothesis is that our finding of robust CPs in CIP, but not in V3A, may be related to the extent to which these areas represent slant in a manner that is tolerant to variations in other cues (e.g., mean disparity). Specifically, it is possible that the lack of CPs in V3A results from a lack of tolerance to changes in mean disparity. We are currently conducting experiments to test this hypothesis directly.

### Dissociating the contributions of stimulus and choice to CIP and V3A activity

To dissociate the contributions of stimulus slant and choice to CIP and V3A responses, we computed partial correlations between these variables and the spike counts of individual neurons. In both areas, we found strong correlations between the stimulus and spike count. In contrast, correlations between choice and spike count were generally strong in CIP, but effectively absent in V3A. This analysis validates the main CP finding; namely, there is strong choice-related activity in CIP but not V3A. These results are reminiscent of a previous study which found that V2, but not V1, neurons show significant choice-related activity during a disparity discrimination task (Nienborg and Cumming, 2006), despite the areas having similar disparity sensitivity. Thus, one potential explanation for these findings is that CPs observed in V2/CIP arise primarily from top-down signals that do not propagate back as strongly to V1/V3A. Another possibility, which is not mutually exclusive, is that the structure of correlated noise is different between V2/CIP and V1/V3A, reflecting that the appearance of CPs may depend on correlated noise (Shadlen et al., 1996; Nienborg and Cumming, 2006; Haefner et al., 2013) and perhaps particularly depend on correlated noise that is information-limiting for the task at hand (Pitkow et al., 2015). An additional possibility, as noted above, is that CIP contains a more invariant representation of slant than V3A.

The pattern of slant and choice partial correlations observed in CIP may reflect a substantial top-down contribution to CPs. In a feedforward (bottom-up) scheme, it would be expected that stimulus and choice partial correlations would have the same sign, such that greater activity from a neuron constitutes evidence in favor of its preferred stimulus. In contrast, our CIP data show no significant relationship between slant and choice partial correlations (Fig. 7A). In other words, slant and choice signals are largely dissociated in CIP, similar to heading and choice signals in VIP (Zaidel et al., 2017). This dissociation may result from top-down choice-related signals that do not target CIP neurons according to their stimulus preferences.

We lastly considered the relative timing of slant and choice signals. The time courses of slant-related signals in CIP and V3A were highly correlated with population-level spike density functions (Fig. 8). In CIP, the time courses of slant-related and choice-related signals differed substantially. Whereas the time course of the slant-related signals peaked around 150-200 ms after stimulus onset and then fell off, the choice-related signals did not become significant until around 350 ms, reached a plateau around 500 ms, and remained elevated until the end of the stimulus. This relatively late choice-related activity in CIP may be consistent with a top-down origin of choice signals in CIP, as suggested above based on the lack of correlation between slant and choice signals. Together, the present findings implicate both V3A and CIP in 3D orientation processing, and reveal a qualitative distinction between the areas since only CIP shows choice-related activity during a fine slant discrimination task.

**Author Contributions**
AR, GCD, and DE Designed research; LCE Performed research; LCE Analyzed data; LCE, AR, GCD, and DE Wrote the paper.

## Acknowledgments

The authors declare no competing financial interests.

This research was supported by NIH grant R01-EY022538 (D.E.A.). L.C.E. was supported by NIH grant F32-EY024515. A.R. was supported by NIH grants R03-DC014305 and R01-EY029438, as well as Whitehall Foundation Research Grant 2016-08-18. G.C.D. was supported by NIH grant R01-EY013644.

## REFERENCES

Alizadeh AM, Van Dromme I, Verhoef BE, Janssen P (2018) Caudal Intraparietal Sulcus and three-dimensional vision: A combined functional magnetic resonance imaging and single-cell study. Neuroimage 166:46–59.

Anzai A, Chowdhury SA, DeAngelis GC (2011) Coding of stereoscopic depth information in visual areas V3 and V3A. J Neurosci 31:10270–10282.

Ban H, Welchman AE (2015) fMRI Analysis-by-Synthesis Reveals a Dorsal Hierarchy That Extracts Surface Slant. J Neurosci 35:9823–9835.

Britten KH, Newsome WT, Shadlen MN, Celebrini S, Movshon JA (1996) A relationship between behavioral choice and the visual responses of neurons in macaque MT. Vis Neurosci 13:87–100.

Celebrini S, Newsome WT (1994) Neuronal and psychophysical sensitivity to motion signals in extrastriate area MST of the macaque monkey. J Neurosci 14:4109–4124.

Chen A, Deangelis GC, Angelaki DE (2013) Functional specializations of the ventral intraparietal area for multisensory heading discrimination. J Neurosci 33:3567–3581.

Chen A, Gu Y, Liu S, DeAngelis GC, Angelaki DE (2016) Evidence for a Causal Contribution of Macaque Vestibular, But Not Intraparietal, Cortex to Heading Perception. J Neurosci 36:3789–3798.

Cumming BG, Nienborg H (2016) Feedforward and feedback sources of choice probability in neural population responses. Curr Opin Neurobiol 37:126–132.

Dodd JV, Krug K, Cumming BG, Parker AJ (2001) Perceptually bistable three-dimensional figures evoke high choice probabilities in cortical area MT. J Neurosci 21:4809–4821.

Durand JB, Nelissen K, Joly O, Wardak C, Todd JT, Norman JF, Janssen P, Vanduffel W, Orban GA (2007) Anterior regions of monkey parietal cortex process visual 3D shape. Neuron 55:493–505.

Galletti C, Battaglini PP (1989) Gaze-dependent visual neurons in area V3A of monkey prestriate cortex. J Neurosci 9:1112–1125.

Goncalves NR, Ban H, Sanchez-Panchuelo RM, Francis ST, Schluppeck D, Welchman AE (2015) 7 tesla FMRI reveals systematic functional organization for binocular disparity in dorsal visual cortex. J Neurosci 35:3056–3072.

Gu Y, DeAngelis GC, Angelaki DE (2007) A functional link between area MSTd and heading perception based on vestibular signals. Nat Neurosci 10:1038–1047.

Gu Y, Angelaki DE, Deangelis GC (2008) Neural correlates of multisensory cue integration in macaque MSTd. Nat Neurosci 11:1201–1210.

Gu Y, Deangelis GC, Angelaki DE (2012) Causal links between dorsal medial superior temporal area neurons and multisensory heading perception. J Neurosci 32:2299–2313.

Haefner RM, Gerwinn S, Macke JH, Bethge M (2013) Inferring decoding strategies from choice probabilities in the presence of correlated variability. Nat Neurosci 16:235–242.

Hillis JM, Watt SJ, Landy MS, Banks MS (2004) Slant from texture and disparity cues: optimal cue combination. J Vis 4:967–992.

Hinkle DA, Connor CE (2002) Three-dimensional orientation tuning in macaque area V4. Nat Neurosci 5:665–670.

Kang I, Maunsell JH (2012) Potential confounds in estimating trial-to-trial correlations between neuronal response and behavior using choice probabilities. J Neurophysiol 108:3403–3415.

Katz LN, Yates JL, Pillow JW, Huk AC (2016) Dissociated functional significance of decision-related activity in the primate dorsal stream. Nature 535:285–288.

Kwon SE, Yang H, Minamisawa G, O’Connor DH (2016) Sensory and decision-related activity propagate in a cortical feedback loop during touch perception. Nat Neurosci 19:1243–1249.

Lewis JW, Van Essen DC (2000) Mapping of architectonic subdivisions in the macaque monkey, with emphasis on parieto-occipital cortex. J Comp Neurol 428:79–111.

Liu Y, Vogels R, Orban GA (2004) Convergence of depth from texture and depth from disparity in macaque inferior temporal cortex. J Neurosci 24:3795–3800.

Moreno-Bote R, Beck J, Kanitscheider I, Pitkow X, Latham P, Pouget A (2014) Information-limiting correlations. Nat Neurosci 17:1410–1417.

Murata A, Gallese V, Luppino G, Kaseda M, Sakata H (2000) Selectivity for the shape, size, and orientation of objects for grasping in neurons of monkey parietal area AIP. J Neurophysiol 83:2580–2601.

Nakamura H, Kuroda T, Wakita M, Kusunoki M, Kato A, Mikami A, Sakata H, Itoh K (2001) From three-dimensional space vision to prehensile hand movements: The lateral intraparietal area links the area V3A and the anterior intraparietal area in macaques. J Neurosci 21:8174–8187.

Nakamura K, Colby CL (2000) Visual, saccade-related, and cognitive activation of single neurons in monkey extrastriate area V3A. J Neurophysiol 84:677–692.

Nguyenkim Jd, DeAngelis GC (2003) Disparity-based coding of three-dimensional surface orientation by macaque middle temporal neurons. J Neurosci 23:7117–7128.

Nienborg H, Cumming BG (2006) Macaque V2 neurons, but not V1 neurons, show choice-related activity. J Neurosci 26:9567–9578.

Nienborg H, Cumming BG (2009) Decision-related activity in sensory neurons reflects more than a neuron’s causal effect. Nature 459:89–92.

Pitkow X, Liu S, Angelaki DE, DeAngelis GC, Pouget A (2015) How can single sensory neurons predict behavior? Neuron 87:411–423.

Prince SJ, Pointon AD, Cumming BG, Parker AJ (2002) Quantitative analysis of the responses of V1 neurons to horizontal disparity in dynamic random-dot stereograms. J Neurophysiol 87:191–208.

Rosenberg A, Angelaki DE (2014a) Gravity influences the visual representation of object tilt in parietal cortex. J Neurosci 34:14170–14180.

Rosenberg A, Angelaki DE (2014b) Reliability-dependent contributions of visual orientation cues in parietal cortex. Proc Natl Acad Sci U S A 111:18043–18048.

Rosenberg A, Cowan NJ, Angelaki DE (2013) The visual representation of 3D object orientation in parietal cortex. J Neurosci 33:19352–19361.

Sanada TM, Nguyenkim JD, Deangelis GC (2012) Representation of 3-D surface orientation by velocity and disparity gradient cues in area MT. J Neurophysiol 107:2109–2122.

Shadlen MN, Britten KH, Newsome WT, Movshon JA (1996) A computational analysis of the relationship between neuronal and behavioral responses to visual motion. J Neurosci 16:1486–1510.

Sugihara H, Murakami I, Shenoy KV, Andersen RA, Komatsu H (2002) Response of MSTd neurons to simulated 3D orientation of rotating planes. J Neurophysiol 87:273–285.

Taira M, Tsutsui KI, Jiang M, Yara K, Sakata H (2000) Parietal neurons represent surface orientation from the gradient of binocular disparity. J Neurophysiol 83:3140–3146.

Tsao DY, Vanduffel W, Sasaki Y, Fize D, Knutsen TA, Mandeville JB, Wald LL, Dale AM, Rosen BR, Van Essen DC, Livingstone MS, Orban GA, Tootell RB (2003) Stereopsis activates V3A and caudal intraparietal areas in macaques and humans. Neuron 39:555–568.

Tsutsui K, Jiang M, Yara K, Sakata H, Taira M (2001) Integration of perspective and disparity cues in surface-orientation-selective neurons of area CIP. J Neurophysiol 86:2856–2867.

Tummers B (2006) DataThief III. In: http://datathief.org/.

Van Dromme IC, Premereur E, Verhoef B-E, Vanduffel W, Janssen P (2016) Posterior Parietal Cortex Drives Inferotemporal Activations During Three-Dimensional Object Vision. PLOS Biol 14:e1002445.

Van Essen DC, Lewis JW, Drury HA, Hadjikhani N, Tootell RB, Bakircioglu M, Miller MI (2001) Mapping visual cortex in monkeys and humans using surface-based atlases. Vision Res 41:1359–1378.

Wichmann FA, Hill NJ (2001) The psychometric function: I. Fitting, sampling, and goodness of fit. Percept Psychophys 63:1293–1313.

Wimmer K, Compte A, Roxin A, Peixoto D, Renart A, de la Rocha J (2015) Sensory integration dynamics in a hierarchical network explains choice probabilities in cortical area MT. Nat Commun 6:6177.

Yang H, Kwon SE, Severson KS, O’Connor DH (2016) Origins of choice-related activity in mouse somatosensory cortex. Nat Neurosci 19:127–134.

Zaidel A, DeAngelis GC, Angelaki DE (2017) Decoupled choice-driven and stimulus-related activity in parietal neurons may be misrepresented by choice probabilities. Nat Commun 8:715.

Zeki SM (1978a) Uniformity and diversity of structure and function in rhesus monkey prestriate visual cortex. J Physiol 277:273–290.

Zeki SM (1978b) The third visual complex of rhesus monkey prestriate cortex. J Physiol 277:245–272.

Zeki SM (1978c) Functional specialisation in the visual cortex of the rhesus monkey. Nature 274:423–428.

